# Efficient Grammar Compression via RLZ-based RePair

**DOI:** 10.1101/2025.07.22.666196

**Authors:** Rahul Varki, Travis Gagie, Christina Boucher

## Abstract

Among grammar-based compression techniques, RePair is a notable offline encoding scheme known for its simplicity and powerful combinatorial properties, producing compact grammars by repeatedly replacing the most frequent adjacent pairs of symbols, known as bigrams. However, RePair’s memory usage scales poorly with input size, as it loads the entire text into memory. In contrast, Relative Lempel-Ziv (RLZ) parsing offers a scalable and lightweight online encoding scheme that losslessly represents a text in terms of phrases that refer to a reference string, but it often fails to expose deeper structural patterns. We introduce an algorithm that produces a RePair grammar from the RLZ parse of the input, leveraging the strengths of both methods. Our method, `RLZ-RePair`, performs bigram replacements systematically, preserving the integrity of the RLZ phrases throughout the RePair iterations. When the reference is well chosen, our method achieves the same grammar as standard RePair while significantly reducing both memory usage and the number of bigram replacements. In particular, we show that `RLZ-RePair` can reduce memory usage by more than 80% while incurring only a modest runtime increase compared to RePair. To our knowledge, `RLZ-RePair` is one of the first scalable methods that constructs exact RePair grammars, resulting in a grammar-based compressor that is both practical for large datasets and faithful to the theoretical elegance of RePair.

## 1 Introduction

In the 1990s, Nevill-Manning and Witten [25] investigated the use of grammar-based compression — specifically, building a context-free grammar generating only the input string — as a means of discovering hierarchical structure in textual data, including natural language, DNA sequences, and music. They obtained promising results but the algorithms they proposed did not scale well to large datasets, such as those arising from biological and web-scale domains. Even implementations of the most popular grammar-based compressors, such as Larsson and Moffat’s RePair [17], tend to use several times more memory than their input (or run very slowly).

RePair constructs a context-free grammar in Chomsky normal form of its input by repeatedly replacing the most frequent pair of adjacent symbols (bigram) with a non-terminal. It is noticeably similar to Gage’s byte-pair encoding (BPE), which is widely used in natural language processing for subword tokenization [10, 15, 26] — which can be viewed as a kind of small-scale hierarchical pattern discovery — in machine translation and training large language models. Whereas RePair continues merging pairs as long as possible, however, BPE merges only for a specified number of rounds. This results in a trade-off: RePair is less efficient but may find large-scale structure that BPE does not.

Following its original development, RePair has been extended and adapted in various ways [2, 5, 6, 11, 21]. Building on these efforts, Gagie et al. [9] proposed using an rsync-style parsing as a preprocessing step to reduce the memory and time overhead of applying RePair on large and repetitive datasets. By parsing the input into phrases and running RePair separately on the dictionary of distinct phrases and on the parse itself, they were able to compress massive inputs more efficiently. More recently, Kim et al. [14] extended the work of Gagie et al. by introducing Re^2^Pair, which leverages recursive prefix-free parsing to further reduce peak memory usage on highly repetitive datasets.

An intuitive next step would be to revisit the goals of Nevill-Manning and Witten, namely, extracting hierarchical structure from large datasets, using the methods of Gagie et al. and Kim et al. However, these methods do not recover the hierarchical structure that traditional RePair does. The core issue lies in the disconnect between the initial parsing and grammar construction. Both Gagie et al. and Kim et al. begin by parsing the input into phrases, using rsync-based chunking in the former and recursive prefix-free parsing in the latter. RePair is then applied to the dictionary, with phrases delimited by unique separators, resulting in a grammar where each phrase is represented by a non-terminal. Simultaneously, RePair is applied to the parse sequence itself, treated as a sequence of phrase identifiers, yielding a second grammar. The two grammars are then merged by replacing each terminal in the parse grammar with the corresponding non-terminal from the dictionary grammar.

These algorithms are efficient because the dictionary and parse are much smaller than the original text in repetitive datasets. However, it imposes an artificial structure on the final grammar, determined not by the actual frequencies of substrings, but by how the initial parsing divided the input. As a result, RePair does not detect frequent substrings that span phrase boundaries. For example, in the original algorithm, a frequently occurring substring is likely to be assigned a dedicated non-terminal, capturing its structural role in the data. In contrast, the methods of Gagie et al. and Kim et al. may arbitrarily break this substring, making it invisible to frequency-based replacement and thus fragmenting the grammar structure. This structural distortion has significant consequences. It makes the grammar less interpretable, limits its utility in downstream tasks that rely on meaningful phrase boundaries, and invalidates the theoretical results derived for traditional RePair grammars. For example, Mieno et al. [22] showed that RePair produces optimal grammars for Fibonacci strings — a result that does not extend to grammars generated via rsync-preprocessed RePair.

These limitations motivate the need for a new approach: one that preserves the scalability of Gagie et al.’s method while maintaining the structural fidelity of the grammar. In this paper, we present such a method, designed to retain the combinatorial properties of RePair while scaling efficiently to large, highly repetitive inputs. Our approach, which we refer to as `RLZ-RePair`, parses the text with RLZ relative to a reference and extracts bigram frequencies in the phrases. Since phrases are intervals that span the reference, we can perform replacements using memory close to that of the reference itself and require fewer substitutions—yet still recover a grammar structurally equivalent to that produced by RePair. In particular, `RLZ-RePair` begins by assigning frequencies to the bigrams derived from the phrases. At each iteration, the most frequent bigram is replaced with a new non-terminal symbol in both the reference and the parse. Special care is taken when bigrams cross phrase boundaries or partially overlap with a phrase source. Since the parse is defined relative to the reference, most replacements in the reference naturally propagate to the parse, requiring minimal additional work. The grammar and bigram frequencies are updated during each round of replacement to reflect the new structure. The result is the RePair grammar, but constructed using far less memory.

We evaluated `RLZ-RePair` on SARS-CoV-2 and human chromosome 19 sequences [27], comparing it against Navarro’s implementation of RePair [17] and, for completeness, against BigRePair [9], and Re^2^Pair [14] — although, as explained above, the latter two algorithms are not useful for discovering hierarchical structure! Our experiments show grammars generated by BigRePair and Re^2^Pair can have far more rules and take noticeably more space than ones generated by RePair and `RLZ-RePair`. `RLZ-RePair` is publicly available at https://github.com/rvarki/RLZ-RePair.

## 2 Preliminaries

### Basic definitions

A string *T* is a finite sequence of symbols *T* = *T* [1..*n*] = *T* [1] …*T* [*n*] over an alphabet Σ = *c*_1_, …, *c*_*σ*_ whose symbols can be unambiguously ordered. We refer to the cardinality of the alphabet Σ as the number of symbols in Σ. We denote by *ε* the empty string, and the length of *T* as |*T*| . We denote by *T* [*i*..*j*] the substring *T* [*i*] … *T* [*j*] of *T* starting at position *i* and ending at position *j*, with *T* [*i*..*j*] = *ε* if *i > j*. For a string *T* and 1 ≤ *i* ≤ *n, T* [1..*i*] is called the *i*-th prefix of *T*, and *T* [*i*..*n*] is called the *i*-th suffix of *T* .

### Context-free Grammars

A context-free grammar (CFG) is a formal grammar in which production rules can be applied to non-terminal symbols regardless of its context. A CFG is formally defined by a set Σ of terminal symbols, a set *V* of non-terminal symbols, a set *R* of production rules, and a start symbol 𝒮. A terminal symbol *c* is a symbol that appears in the original text, whereas a non-terminal symbol *β* is a new symbol not a part of Σ introduced into the text. A production rule defines how a non-terminal symbol decompresses to a sequence of terminal and non-terminal symbols . The production rules are written in the form of *β* → *α*, where *α* defines a consecutive sequence of *c* and *β* symbols that appear in the text. The start symbol is defined as the initial non-terminal symbol from which the original text can be reconstructed by applying the production rules. In practice, a CFG is defined by its start symbol and production rules, where the sets of terminal and non-terminal symbols are implicitly defined by these rules.

Chomsky normal form is a CFG that requires that all production rules adhere to one of the following forms: (1) *β*_*i*_ → *β*_*j*_*β*_*k*_ or (2) *β*_*i*_ → *c*_*i*_ where *β*_*i*_, *β*_*j*_, *β*_*k*_ ∈ *V* and *i > j, k*, and *c*_*i*_ ∈ Σ. In other words, a non-terminal symbol should either decompress into (1) two other non-terminal symbols or (2) a single terminal symbol.

A straight line program (SLP) is a CFG in Chomsky normal form that derives exactly one string. An SLP is a lossless grammar-based compression scheme that represents an input text *T* . With an SLP, random access for any substring of text can be achieved with an additive logarithmic time penalty [1, 4]. For brevity, we refer to the SLP that produces an input text *T* simply as a *grammar of T* . We denote the compressed representation of *T* in the SLP as *T*.𝒞. Similarly, we denote the set of production rules in the SLP as *T*.ℛ.

### Relative Lempel-Ziv

Relative Lempel-Ziv (RLZ), introduced by Kuruppu et al. [16], is a dictionary-based compression scheme designed to efficiently compress large collections of similar sequences, such as those found in genomic datasets. Unlike traditional Lempel-Ziv variants that parse a sequence based on previous substrings within the same sequence [13, 8], RLZ uses a reference string to guide the parsing of each target sequence. Hence, given a reference string *R* and a target string *T*, the RLZ algorithm parses *T* into a sequence of phrases, where each phrase corresponds to the longest prefix of the remaining suffix of *T* that matches a substring of *R*. Each phrase is encoded as a pair (*p, ℓ*), where *p* denotes the starting position of the matching substring in *R* and *ℓ* denotes the length of the match. If no match is found in *R*, either it can fail or a single character (literal) from *T* can be emitted. The reference string can be uncompressed or compressed with a method that supports fast access [24].

## 3 RLZ-RePair

### 3.1 Overview

`RLZ-RePair` combines RLZ parsing with RePair compression to construct an exact, compact grammar using less memory than the standard RePair algorithm. It applies RePair to the RLZ parse and reference, which are often much smaller than the input, keeping memory usage close to the size of the reference when its well chosen. This makes `RLZ-RePair` particularly effective for large and repetitive inputs. To motivate the algorithm, we begin by introducing an example.

Suppose we want to apply `RLZ-RePair` to the text *T* [1..15] = TGAAACTAAATGCTC using the reference *R*[1..6] = `GAAACT`. We first compute the RLZ parse of *T* with respect to *R*: *T* = {`(6, 1), (1, 6), (2, 3), (6, 1), (1, 1), (5, 2), (5, 1)`} . For convenience, we reinterpret these pairs as endpoints of the intervals they span: *T* = {`(6, 6), (1, 6), (2, 4), (6, 6), (1, 1), (5, 6), (5, 5)`} . After applying RLZ, we then compute bigram frequencies in *T* using the RLZ parse, as shown in Figure 1. Once this information is calculated, we can start to apply the RePair algorithm.

**Figure 1.**
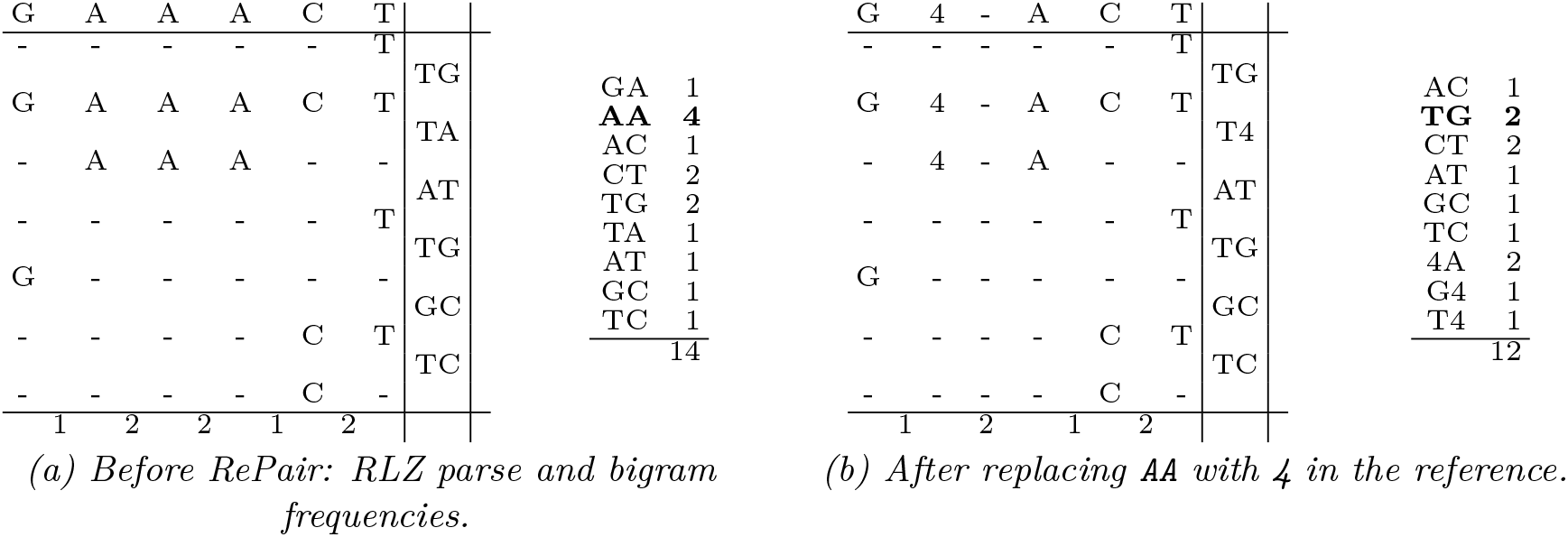
Visual comparison of RLZ parses and derived bigram frequencies before (a) and after (b) replacing `AA` with `4` in the reference. In (a), total bigram frequencies in *T* sum to 14; in (b), they sum to 12.

From the frequency table in Figure 1(a), we find that the bigram `AA` occurs with highest frequency. Before performing the replacement, the algorithm checks whether this would invalidate any of the RLZ phrases. In this case, the figure shows that all occurrences of `AA` in *T* occur entirely within RLZ phrases, with none partially overlapping an occurrence in *R*. This indicates that it is safe to do the replacement. Since all occurrences of `AA` are within RLZ phrases, the algorithm can perform the replacement by simply replacing all instances of the bigram in *R*, as this will naturally propagate to the RLZ phrases that span these occurrences. Figure 1(b) shows the updated information after the replacement. If *T* is repetitive and *R* captures that repetitiveness, the majority of replacements will follow the same pattern. However, in some cases, preprocessing is required before the replacement can occur. These cases are described in the following sections.

### 3.2 RLZ Construction

The algorithm begins with RLZ parsing. Given an input text *T* and a reference text *R, T* is parsed into a sequence of phrases, where each phrase corresponds to the longest prefix of the remaining suffix of *T* that matches a substring of *R*. Each phrase *P*_*i*_ can be represented as a pair (*p*_*i*_, *ℓ*_*i*_), where *p*_*i*_ denotes the starting position of the match in *R*, and *ℓ*_*i*_ denotes its length. Each phrase spans a section of the reference, forming a dictionary-style encoding of *T* . The parse can be denoted as *T* = *P*_1_ |*P*_2_ | … | *P*_*k*_, where each *P*_*i*_ is a phrase derived from *R*. The RLZ parse effectively captures repetitive patterns in the text when an appropriate reference sequence is selected, i.e., when *k* ≪ |*T* |.

### 3.3 RePair Construction

Initially, the reference *R* and RLZ phrases (*P*_1_ |*P*_2_ | … | *P*_*k*_) are loaded into memory. Rather than storing each phrase as a (*p*_*i*_, *ℓ*_*i*_) pair, we represent them as (*s*_*i*_, *e*_*i*_) intervals, where *s*_*i*_ and *e*_*i*_ denote the starting and ending positions in *R* that the phrase spans. We note that references to positions in *R* denote *logical*, not absolute, positions; in the following equations, +1 and -1 indicate the logically adjacent positions. Returning to the discussion of phrase intervals, we note that initially *s*_*i*_ = *p*_*i*_ and *e*_*i*_ = *p*_*i*_ + *ℓ*_*i*_ − 1. Hence, we obtain the following definition.

#### Definition 1

`(Non-Explicit Phrase)`. *Let R* ∈ Σ^∗^ *be the reference string, and let T* ∈ Σ^∗^ *be a string divided into k RLZ phrases* (*P*_1_ |*P*_2_ | … | *P*_*k*_) *with respect to R. Each phrase P*_*i*_ *is originally represented as a tuple* (*p*_*i*_, *ℓ*_*i*_), *where p*_*i*_ *is the starting position in R and ℓ*_*i*_ *is the length of the phrase. We define a* non-explicit phrase *as the interval* (*s*_*i*_, *e*_*i*_), *where at the beginning s*_*i*_ = *p*_*i*_ *and e*_*i*_ = *p*_*i*_ + *ℓ*_*i*_ − 1, *indicating the start and end positions in R that the phrase spans*.

Bigram frequencies are then computed over the set of non-explicit phrases, including those occurring entirely within phrases and those spanning phrase boundaries. This computation is performed only once at the start of the algorithm. The frequencies are maintained throughout the algorithm once they are initially computed. Next, the most frequent bigram is selected for replacement. This can occur entirely within phrases, spanning phrase boundaries, or both. We now describe the behavior of the algorithm in each scenario. This selection and replacement are repeated for each iteration of the algorithm.

#### 3.3.1 Bigram Substitution Within Phrase Boundaries

If the most frequent bigram in *T* also occurs in *R* and is fully covered by non-explicit phrases, then the replacements only need to occur in *R*. Recall from Definition 1 that non-explicit phrases are intervals that span the reference sequence. Replacements in *R* are automatically reflected in the non-explicit phrases that reference it, leading to the following observation.

▶ **Observation 2**. *Let R* ∈ Σ^∗^ *be the reference string and let R[i]R[i+1] be an occurrence of the most frequent bigram in T* . *By Definition 1, a non-explicit phrase refers to an interval in the reference. If there exist n non-explicit phrases that completely span the interval (i,i+1), then the number of required replacements is reduced by n-1*.

Additionally, if we restrict our attention to non-explicit phrases in *R* that strictly contain the interval, meaning that the interval lies entirely within the phrase and does not touch its boundaries, then the update process becomes simpler. In this case, updating the bigram frequencies requires only four operations: two decrements to reduce the counts of the replaced bigrams, and two increments to increase the counts of the new bigrams. However, if the bigram occurrence in *R* touches the boundary of a non-explicit phrase, updating its frequencies requires examining the boundary character of its adjacent phrase.

A potential complication of replacing a bigram in *R* is that it shortens *R*, which would invalidate the intervals of the non-explicit phrases. To avoid this, *R* can be stored as a doubly linked list embedded in an array (see Figure 2). This not only allows fast random access within *R*, but also ensures that when a character is deleted as a result of a merge, only the pointers of adjacent nodes need to be updated to bypass it, thereby preserving the intervals of the phrases. However, because *R* is not shortened, adjacent non-deleted positions are not necessarily adjacent in absolute coordinates. This is the reason why we refer to the positions in *R* in logical rather than absolute terms.

**Figure 2.**
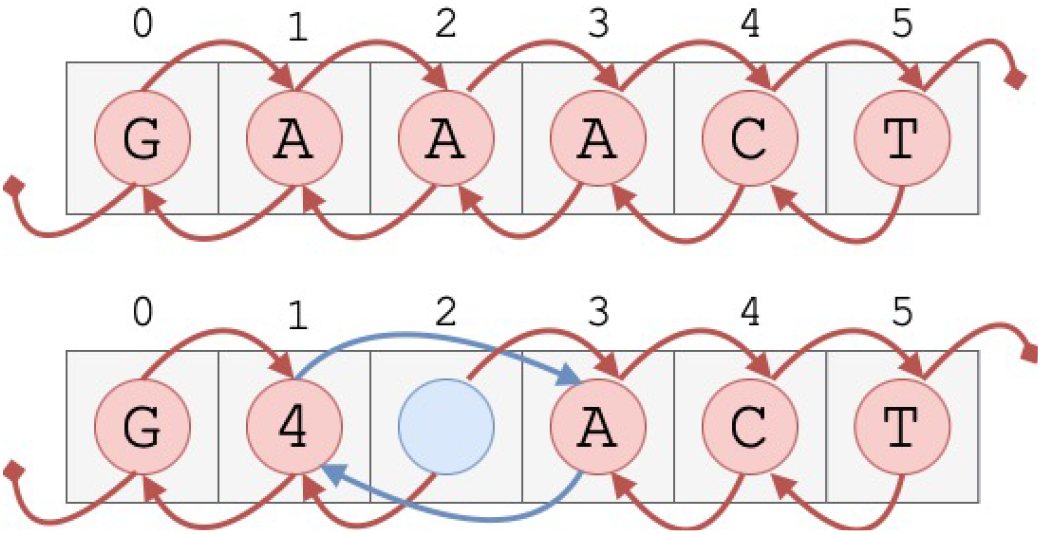
Reference represented as a linked list embedded within an array. The top diagram shows the reference before any replacements for the example introduced in the RLZ-RePair Overview section. The bottom diagram shows the reference after replacing all occurrences of `AA` with the new symbol `4`. Although `AA` occurs twice in the reference, only one replacement is needed since the occurrences are consecutive.

A carefully selected reference *R* that effectively captures the repetitions in *T* yields a small number of long non-explicit phrases. Consequently, most replacements occur within phrase boundaries, substantially reducing the total number of replacements required. This characteristic represents one of the key advantages of the algorithm.

#### 3.3.2 Bigram Substitution Under Boundary Constraints

For the following explanation, we let *X* = *ab* denote the most frequent bigram identified in *T* . Before substituting all occurrences of *X* in *R*, the algorithm ensures that the replacement will not invalidate any non-explicit phrase. A non-explicit phrase (*s*_*i*_, *e*_*i*_) will become invalid under two conditions:

1. The bigram *X* occurs across the boundary between two consecutive phrases *P*_*i*_ and *P*_*i*+1_, i.e., the character *R*[*e*_*i*_] = *a* and *R*[*s*_*i*+1_] = *b*.
2. The bigram *X* occurs in *R* such that it partially overlaps with the interval [*s*_*i*_, *e*_*i*_], i.e., either *s*_*i*_ = *j* + 1 or *e*_*i*_ = *j* for some occurrence of *ab* = *R*[*j*]*R*[*j* + 1] in the reference.

Since non-explicit phrases must exactly match substrings of the reference *R*, these scenarios violate the constraint that *R*[*s*_*i*_, *e*_*i*_] defines a valid interval in *R*. To restore this invariant, the boundaries of the affected non-explicit phrase (*s*_*i*_, *e*_*i*_) are adjusted as follows:

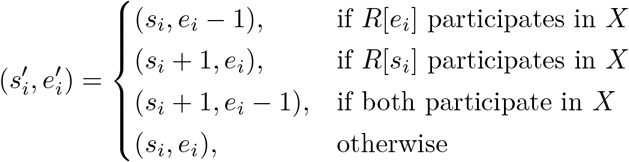

The character(s) removed, i.e., *R*[*e*_*i*_] and / or *R*[*s*_*i*_], are stored explicitly as literals in a newly created explicit phrase *E*, which is inserted adjacently to the affected phrases. We formalize this with the following definition.

##### Definition 3

`(Explicit Phrase)`. *An* explicit phrase *is a sequence of uncompressed boundary characters introduced during an iteration. These characters originate from adjacent non-explicit phrases that became invalid*.

We now examine each of these scenarios in detail and describe the corrective transformations required to preserve the structural correctness of the non-explicit phrases.

#### 3.3.3 Phrase Boundary Condition

If the most frequent bigram spans two consecutive phrases, this is called a *phrase boundary condition*. When this occurs and the two consecutive phrases are non-explicit phrases *P*_*i*_ and *P*_*i*+1_, the characters corresponding to *e*_*i*_ and *s*_*i*+1_ are removed from their respective phrases and stored explicitly in a new explicit phrase positioned between the phrases, i.e., *P*_*i*_ | *E* | *P*_*i*+1_. Let 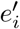 and 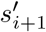 be the new end and start position of the non-explicit phrases after the removal, then 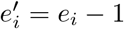 and 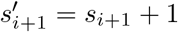. If one of the phrases involved in the phrase boundary condition is already explicit, only the non-explicit phrase will lose one boundary character, which will then be added either to the beginning (i.e., *P*_*i*_ | *E*) or end (i.e., *E* | *P*_*i*_) of the explicit phrase. In addition to the fact that the bigram replacement cannot directly occur if it spans two phrases, this condition invalidates the non-explicit phrases because the phrases are guaranteed to be non-consecutive in relation to *R*; otherwise, they would have been merged initially. In other words, since this replacement does not exist in *R*, the bigram must be made explicit prior to replacement.

#### 3.3.4 Source Boundary Condition

If a non-explicit phrase’s start or end position in *R* partially overlaps with an occurrence of the bigram to be replaced, this is called a *source boundary condition*. For a non-explicit phrase *P*_*i*_, if its starting position *s*_*i*_ partially overlaps an occurrence of the bigram in *R*, then the characters at positions *s*_*i*_ − 1 and *s*_*i*_ in *R* form the bigram. In this case, a new explicit phrase is formed to the left of the non-explicit phrase, unless an explicit phrase already exists there (i.e., *E* | *P*_*i*_). The character at position *s*_*i*_ is made explicit and added to the *end* of the explicit phrase, and the new start position of the non-explicit phrase is 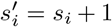. Similarly, if its end position *e*_*i*_ partially overlaps with an occurrence of the bigram in *R*, then that means that the characters at positions *e*_*i*_ and *e*_*i*_ + 1 form the bigram in *R*. When this occurs, a new explicit phrase is formed to the right of the non-explicit phrase, unless an explicit phrase already exists there (i.e., *P*_*i*_ | *E*). The character at position *e*_*i*_ is made explicit and added to *start* of the explicit phrase, and the new end position of the non-explicit phrase is 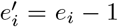. This condition invalidates non-explicit phrases because replacing the bigram occurrence in *R* can cause the boundary characters of some non-explicit phrases to reference non-existent (deleted) positions in the reference. Therefore, these boundary characters must be made explicit prior to the replacement in order to preserve the integrity of the non-explicit phrases.

After making all necessary characters explicit, the algorithm proceeds with the bigram replacements in the phrases. As explained earlier, for non-explicit phrases, replacements only need to occur in *R*, as they all refer to it, a primary benefit of the algorithm. If *R* is significantly smaller than the input, then the bigram is likely to occur less often in it. Therefore, if non-explicit phrases cover most of the input, then the number of replacements needed is typically much smaller than in standard RePair. For explicit phrases, the algorithm performs the replacement similar to standard RePair.

### 3.4 Implementation

The algorithm relies on several auxiliary data structures to enable efficient processing and updates. A max-heap tracks the frequencies of all bigrams, including those within and across phrase boundaries. An augmented dynamic implicit interval tree maintains intervals spanned by non-explicit phrases, enabling efficient identification of phrases that completely contain an interval. This tree is an augmented Red-Black Tree where each node stores its interval, as well as the minimum and maximum values of its subtree—unlike traditional implicit interval trees, which typically store only the interval and maximum value [20, 7]. A hash table maps each phrase to the bigram that extends its boundary, supporting efficient identification of phrase boundary bigrams. Additional hash tables map the first and last characters of non-explicit phrases, enabling quick identification of potential source boundary-crossing bigrams. Lastly, a hash table maps each bigram in explicit phrases to allow for direct access during replacement.

## 4 Experiments

We evaluated the performance of our method, `RLZ-RePair` in compressing sequences from two biologically distinct datasets: 400,000 SARS-CoV-2 genomes (viral) and 1,024 human chromosome 19 assemblies (mammal) [27]. For both datasets, we progressively compressed larger subsets of the full dataset. Since RLZ parsing is sensitive to the choice of reference file, we tested `RLZ-RePair` with multiple reference files for each dataset. Specifically, we randomly subsampled 0.5%, 1%, and 2% of the sequences in the full dataset to use as references, where each subsampled reference was a superset of the previous. We randomly subsampled from the full FASTA file using seqtk (v1.3-r106) [18] with the command: `seqtk sample -s 100 [FASTA] [fraction]`, where `[fraction]` was set to 0.005, 0.01, or 0.02. We denote different software configurations by placing the configuration name in brackets after the software name. We compared the performance of `RLZ-RePair` with Navarro’s version of standard RePair [17] (http://www.dcc.uchile.cl/gnavarro/software/repair.tgz). Due to `malloc` errors encountered when running the default version of RePair on larger collections, we also evaluated the large memory variant which produces balanced grammars to ensure compatibility with larger inputs. Furthermore, we compared our performance against BigRePair [9] and Recursive RePair (Re^2^Pair) [14], methods that produce RePair-style grammars. Our results show that `RLZ-RePair` achieves significant memory savings over RePair, with only a modest increase in runtime.

`RLZ-RePair` was implemented using C++ and Python, with the max heap and decompression code borrowed from the C implementation of Navarro’s version of RePair. The code was compiled using CMake (v3.30.5) with GCC (v12.2.0). The experiments were carried out on a server with 100 GB RAM and an AMD EPYC 75F3 32-core CPU clocked at 2.95 GHz, running Python (v3.11). The wall clock time and maximum memory usage were measured for each tool using Snakemake (v7.32.4)[23], running all tools on a single thread. Experiments exceeding 100 GB of RAM or 24 hours were omitted from further analysis.

### 4.1 Results on SARS-CoV-2

For the SARS-CoV-2 dataset, we compressed subsets of 25,000, 50,000, 100,000, 200,000, and 400,000 genomes, each larger subset being a superset of the previous. To provide context, the 25,000 subset contained 744,740,774 characters (0.74 GB) while the 400,000 subset contained 11,931,360,555 characters (11.93 GB). We present the RLZ statistics produced by each `RLZ-RePair` configuration for the entire 400,000 dataset in Table 1. The table shows that as the number of reference sequences increased, the number of RLZ phrases decreased, while both the average phrase length and its standard deviation increased proportionally. This suggests that the dataset is highly repetitive and that the observed trends were largely driven by the inclusion of more input sequences in the reference. The high similarity between SARS-CoV-2 sequences has been observed by others [12, 19]. We note that the observed trends were consistent across the other subsets.

**Table 1.** RLZ statistics for the SARS-CoV-2 400K dataset. From left to right, the columns show the number of characters in the reference (*n*_*ref*_), number of sequences in the reference (*m*_*ref*_), number of RLZ phrases (No. Phrases), average RLZ phrase length (Mean), and standard deviation of phrase length (SD).

| Tool | $n_{ref}$ | $m_{ref}$ | No. Phrases | Mean | SD |
| --- | --- | --- | --- | --- | --- |
| RLZ-RePair (0.5%) | 59,151,937 | 1,983 | 5,084,916 | 2,346 | 4,219 |
| RLZ-RePair (1%) | 119,346,706 | 4,001 | 4,565,438 | 2,613 | 4,612 |
| RLZ-RePair (2%) | 236,727,396 | 7,936 | 4,093,433 | 2,914 | 5,086 |

We show in Figure 3 the compression wall clock time and peak memory usage of all tools across all subsets. As the number of sequences in the collection increased, we observed that all configurations of `RLZ-RePair` used significantly less peak memory than RePair, with only a modest decrease in compression speeds. To compress the full set of 400,000 SARS-CoV-2 sequences, `RLZ-RePair` (0.5%), the most memory efficient configuration, used 17.17 GB and took 4,942 seconds. Compared to RePair (large_bal), which used 99.88 GB and took 3,875 seconds, `RLZ-RePair` (0.5%) required 82.8% less memory while only being 27.5% slower. For context, `RLZ-RePair` (2%), the least memory efficient configuration, used 18.72 GB and took 4,804 seconds, only 1.55 GB more and 138 seconds faster than `RLZ-RePair` (0.5%). We were unable to get RePair (default) to run past 50,000 sequences, but it used approximately half the memory of RePair (large_bal). In general, the most memory efficient and fastest methods in this experiment were BigRePair and Re^2^Pair, with the former using 1.98 GB and taking 357 seconds, and the latter using only 2.21 GB and 244 seconds.

**Figure 3.**
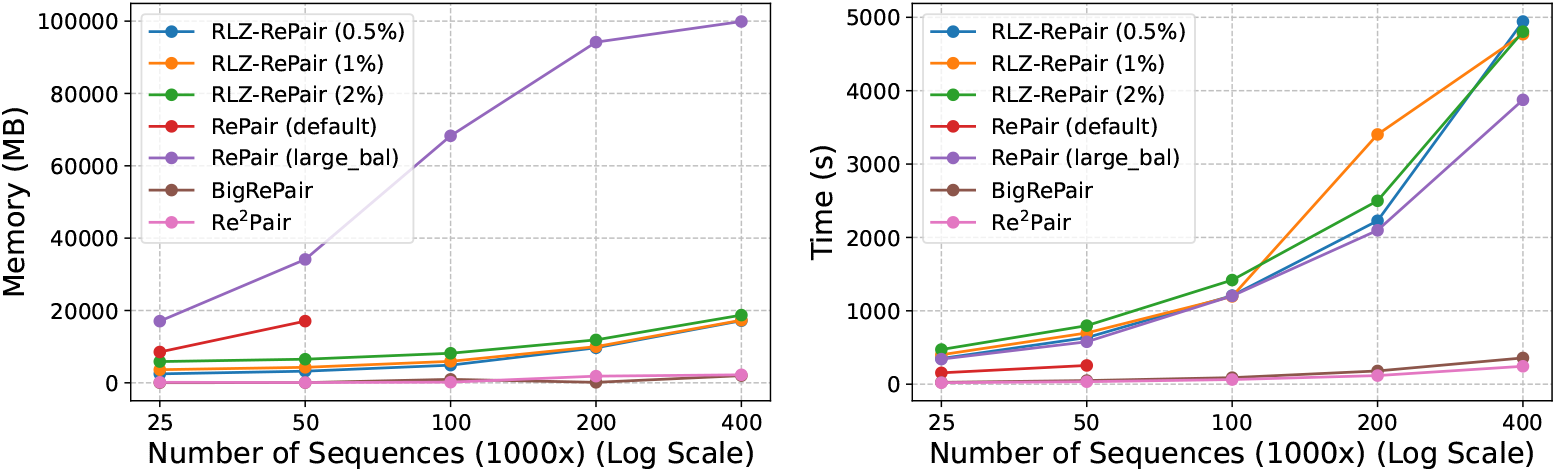
Resource usage in SARS-CoV-2 compression. The left figure shows the peak memory usage (MB), and the right figure shows the wall-clock time (s) required by each tool to compress the subsets.

When comparing the performance trends of `RLZ-RePair` and RePair, we observe that all configurations of `RLZ-RePair` and RePair exhibit linear scaling in both runtime and memory usage. Notably, RePair (large_bal) begins to show sublinear memory growth beyond 100,000 sequences—an artifact of nearing system memory limits—which ultimately leads to memory thrashing at larger scales (e.g., 400,000 sequences). In contrast, `RLZ-RePair` maintains stable and predictable resource usage throughout. As shown in Table 2, all configurations of `RLZ-RePair` achieve compression ratios identical to those of RePair (large_bal) on the full SARS-CoV-2 dataset. The compressed file sizes and number of production rules are nearly indistinguishable, with minor differences attributable to implementation-level details that do not affect correctness. This consistency is preserved across all evaluated dataset sizes. Importantly, both `RLZ-RePair` and RePair yield more compact grammars than BigRePair and Re^2^Pair, which generate shorter compressed sequences only by introducing a significantly larger number of rules. Specifically, `RLZ-RePair` and RePair compressed the full 11.93 GB file to 20.48 MB, compared with 24.64 MB (20% larger) for BigRePair and 34.88 MB (70% larger) for Re^2^Pair. Since these alternatives do not construct exact RePair grammars, they cannot guarantee the same structural fidelity. In contrast, `RLZ-RePair` preserves the full semantics and guarantees of the original RePair algorithm while offering strong scalability and memory efficiency—making it a compelling choice for large-scale grammar-based compression.

**Table 2.**
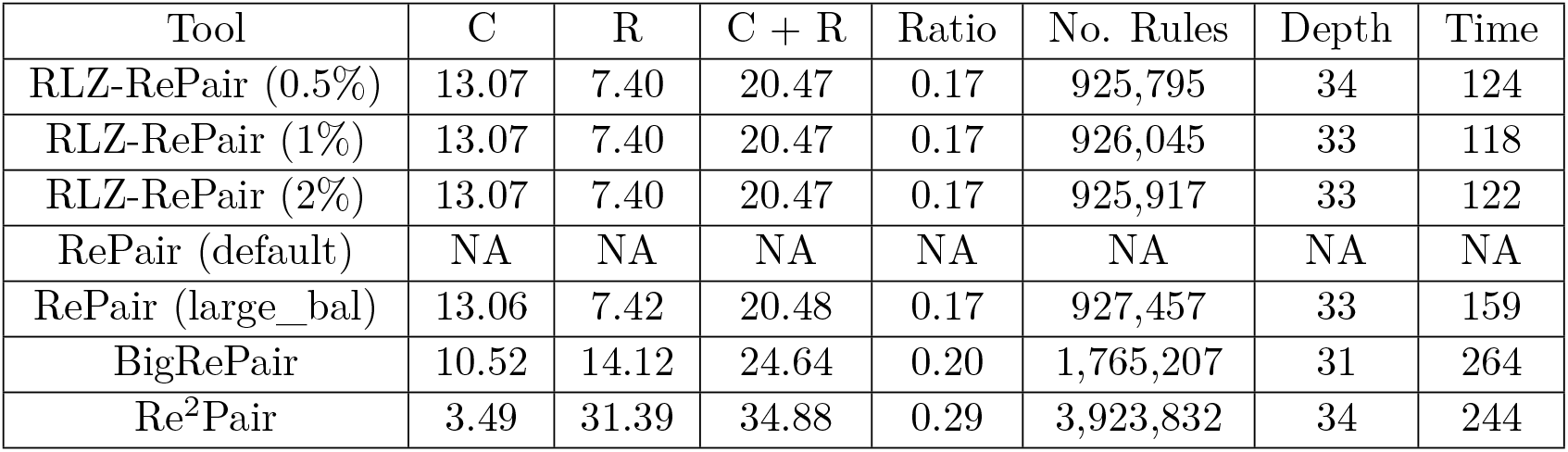
Compression results for 400,000 SARS-CoV-2 sequences. Reported are the compressed size (C), rule size (R), total size (C + R), and compression ratio (%). Additionally, the number of rules (No. Rules), max rule depth (Depth), and decompression time in seconds (Time) are reported. All size values are reported in megabytes (MB). The uncompressed size is 11,931.36 MB. The compression ratio is calculated as total size divided by uncompressed size times 100. NA is reported when a tool fails to run to completion.

| Tool | C | R | C + R | Ratio | No. Rules | Depth | Time |
| --- | --- | --- | --- | --- | --- | --- | --- |
| RLZ-RePair (0.5%) | 13.07 | 7.40 | 20.47 | 0.17 | 925,795 | 34 | 124 |
| RLZ-RePair (1%) | 13.07 | 7.40 | 20.47 | 0.17 | 926,045 | 33 | 118 |
| RLZ-RePair (2%) | 13.07 | 7.40 | 20.47 | 0.17 | 925,917 | 33 | 122 |
| RePair (default) | NA | NA | NA | NA | NA | NA | NA |
| RePair (large_bal) | 13.06 | 7.42 | 20.48 | 0.17 | 927,457 | 33 | 159 |
| BigRePair | 10.52 | 14.12 | 24.64 | 0.20 | 1,765,207 | 31 | 264 |
| Re<sup>2</sup>Pair | 3.49 | 31.39 | 34.88 | 0.29 | 3,923,832 | 34 | 244 |

### 4.2 Results on Chromosome 19

Lastly, we benchmarked the performance of `RLZ-RePair` by compressing subsets of 1, 2, 4, 8, 16, 32, 64, 128, 256, 512, and 1024 sequences of human chromosome 19 (chr19), each larger subset being a superset of the previous. For context, the single copy of chr19 contained 59,128,983 characters (0.059 GB), while the full set of 1,024 chr19 sequences contained 60,543,372,726 characters (60.54 GB). Table 3 shows the RLZ statistics for each `RLZ-RePair` configuration on the entire 1,024 dataset. The table shows that as the number of reference sequences increased, the number of RLZ phrases decreased, while both the average phrase length and its standard deviation increased. Similar to the SARS-CoV-2 dataset, this suggests that the dataset is repetitive and that the observed trends were largely driven by including more input sequences in the reference.

**Table 3.** RLZ statistics for the 1,024 chromosome 19 dataset. From left to right, the columns show the number of characters in the reference (*n*_*ref*_), number of sequences in the reference (*m*_*ref*_), number of RLZ phrases (No. Phrases), average RLZ phrase length (Mean), and standard deviation of phrase length (SD).

| Tool | $n_{ref}$ | $m_{ref}$ | No. Phrases | Mean | SD |
| --- | --- | --- | --- | --- | --- |
| RLZ-RePair (0.5%) | 472,993,586 | 8 | 20,900,644 | 2,896 | 43,146 |
| RLZ-RePair (1%) | 886,865,353 | 15 | 13,668,505 | 4,429 | 68,206 |
| RLZ-RePair (2%) | 1,300,736,553 | 22 | 10,074,413 | 6,009 | 97,148 |

We show in Figure 4 the compression wall clock time and peak memory usage of all tools in all subsets. For collections larger than 32 sequences, all configurations of `RLZ-RePair` used significantly less peak memory than RePair (large_bal) while only being moderately slower. However, neither RePair (default) nor RePair (large_bal) could compress the full chr19 dataset within the given resource constraints. The largest collection RePair (large_bal) could handle was 256 sequences. It exceeded the 24-hour time limit when compressing 512 sequences, presumably due to thrashing after reaching the 100 GB memory limit. In contrast, across all configurations, `RLZ-RePair` used between 31.09 GB and 41.62 GB to compress all 1,024 sequences, handling four times as many sequences while using less than half the available memory.

**Figure 4.**
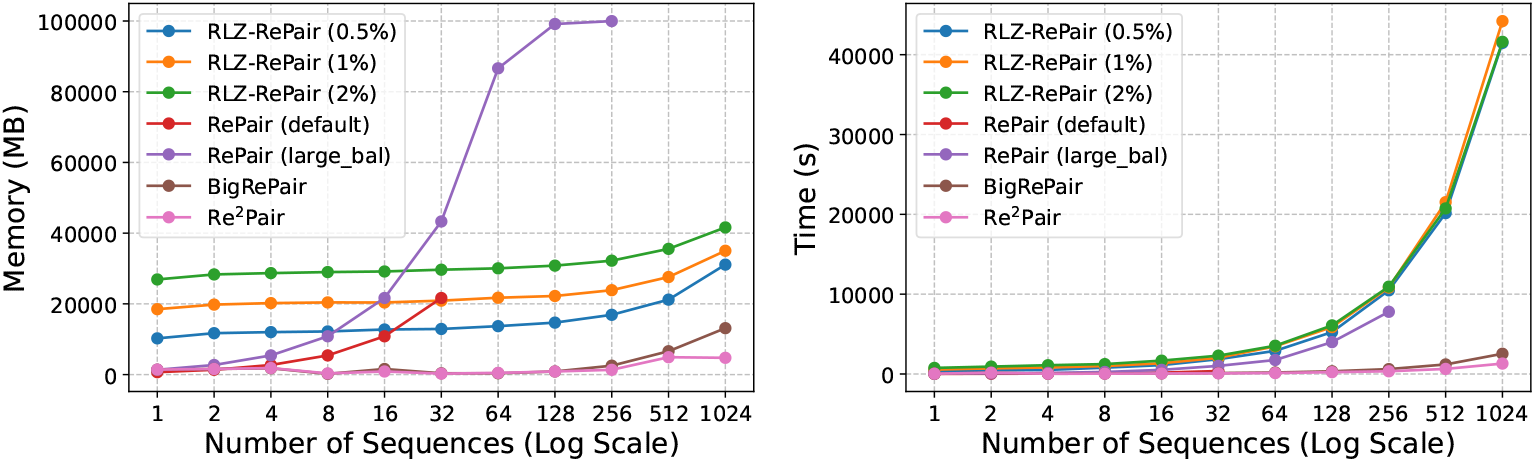
Resource usage in chromosome 19 compression. The left figure shows the peak memory usage (MB), and the right figure shows the wall-clock time (s) required by each tool to compress the subsets.

To evaluate relative performance against RePair, we consider the 256-sequence subset, the largest RePair (large_bal) could compress. For 256 sequences, `RLZ-RePair` (0.5%), the most memory efficient configuration, used 16.90 GB and took 10,480 seconds, whereas RePair (large_bal) used 100 GB and took 7,793 seconds. Thus, `RLZ-RePair` (0.5%) used 83.1% less memory while being 34.5% slower compared to RePair (large_bal). For context, `RLZ-RePair` (2%), the least memory efficient configuration, used 32.21 GB and took 10,942 seconds, 15.31 GB more and 462 seconds slower than `RLZ-RePair` (0.5%). We again observe that the most memory and time efficient methods were BigRePair and Re^2^Pair, with the former using 2.51 GB and taking 614 seconds, and the latter using 1.34 GB and taking 340 seconds.

As in the previous experiment, Table 4 shows that all configurations of `RLZ-RePair` produced grammars approximately the same size as RePair (large_bal) for the 256 sequences, a trend consistent across all smaller subsets. The grammars were consistently smaller than those generated by BigRePair and Re^2^Pair. Specifically, `RLZ-RePair` and RePair compressed the 15.13 GB file to 69.02 MB compared with 74.38 MB (8% larger) for BigRePair and 78.89 MB (14% larger) for Re^2^Pair. Table 5 shows the grammar statistics for the 1,024 sequences. Since no RePair configuration could compress the full dataset, we could not confirm whether `RLZ-RePair` produced a grammar of the same size, though it is likely. However, we do observe that `RLZ-RePair` compressed the 60.54 GB file to 107.90 MB compared with 118.11 MB (10% larger) for BigRePair and 131.61 MB (22% larger) for Re^2^Pair.

**Table 4.**
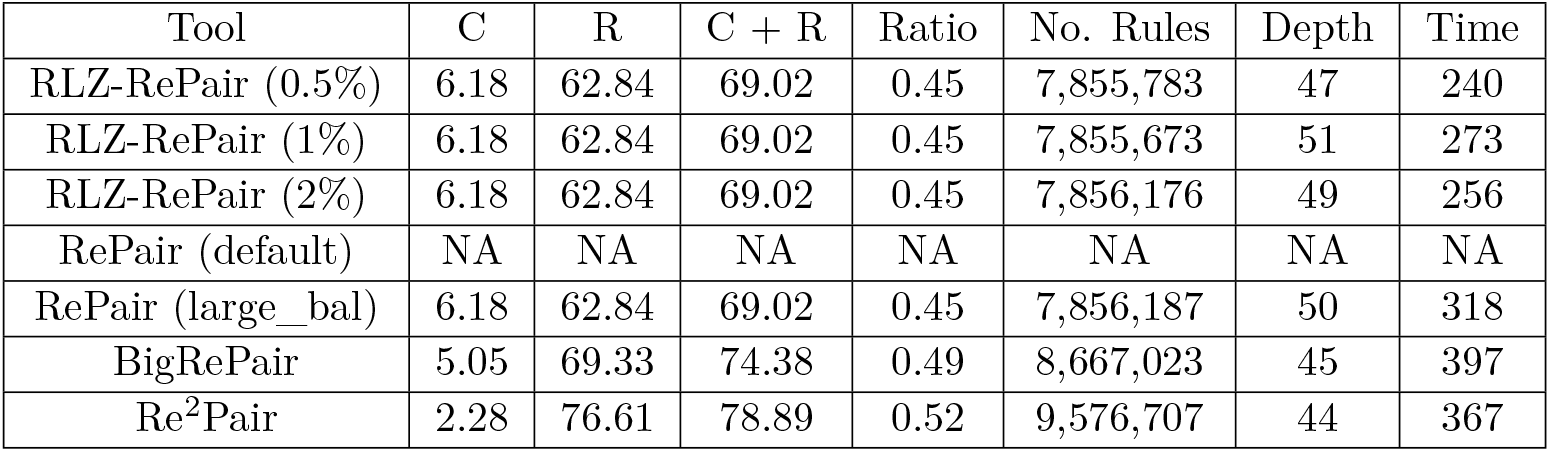
Compression results for 256 chromosome 19 sequences. Reported are the compressed size (C), rule size (R), total size (C + R), and compression ratio (%). Additionally, the number of rules (No. Rules), max rule depth (Depth), and decompression time in seconds (Time) are reported. All size values are reported in megabytes (MB). The uncompressed size is 15,135.85 MB. The compression ratio is calculated as total size divided by uncompressed size times 100. NA is reported when a tool fails to run to completion.

**Table 5.**
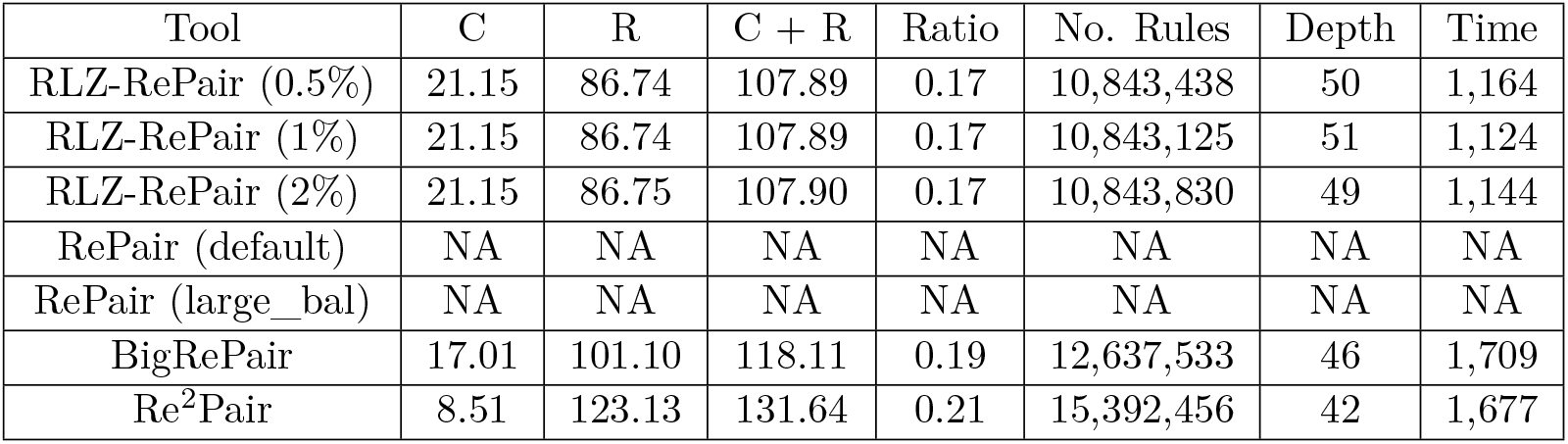
Compression results for 1,024 chromosome 19 sequences. Reported are the compressed size (C), rule size (R), total size (C + R), and compression ratio (%). Additionally, the number of rules (No. Rules), max rule depth (Depth), and decompression time in seconds (Time) are reported. All size values are reported in megabytes (MB). The uncompressed size is 60,543.37 MB. The compression ratio is calculated as total size divided by uncompressed size times 100. NA is reported when a tool fails to run to completion.

| Tool | C | R | C + R | Ratio | No. Rules | Depth | Time |
| --- | --- | --- | --- | --- | --- | --- | --- |
| RLZ-RePair (0.5%) | 21.15 | 86.74 | 107.89 | 0.17 | 10,843,438 | 50 | 1,164 |
| RLZ-RePair (1%) | 21.15 | 86.74 | 107.89 | 0.17 | 10,843,125 | 51 | 1,124 |
| RLZ-RePair (2%) | 21.15 | 86.75 | 107.90 | 0.17 | 10,843,830 | 49 | 1,144 |
| RePair (default) | NA | NA | NA | NA | NA | NA | NA |
| RePair (large_bal) | NA | NA | NA | NA | NA | NA | NA |
| BigRePair | 17.01 | 101.10 | 118.11 | 0.19 | 12,637,533 | 46 | 1,709 |
| Re<sup>2</sup>Pair | 8.51 | 123.13 | 131.64 | 0.21 | 15,392,456 | 42 | 1,677 |

## Conclusion

We introduced `RLZ-RePair`, a method for RePair compression via RLZ parsing that avoids loading the input text into memory and significantly reduces replacements while preserving the exact RePair grammar. Unlike BigRePair and Re^2^Pair, which sacrifice RePair’s theoretical guarantees for speed and memory efficiency, `RLZ-RePair` reduces memory usage while preserving them. Although `RLZ-RePair` is designed to produce grammars equivalent to those of the original RePair, our implementation may yield slight variations compared to Navarro’s RePair due to minor algorithmic differences. First, when a tie occurs, our method chooses the most frequent bigram using the same max heap structure as Navarro’s RePair, which aligns with the original RePair algorithm described by Larson and Moffat [17]. In their paper, they acknowledge this scenario and consider it of “minor importance”, opting for the least recently accessed maximally frequent bigram for replacement. Since our implementation performs replacements in a non-deterministic order using unordered hash tables to find the occurrences, the least accessed maximally frequent bigram may vary between runs, leading to slightly different grammars. The second difference is the choice of bigram to replace when there are consecutive runs of the same symbol. For example, if the bigram is CC and CCC appears in the text, Navarro’s RePair replaces the last occurrence (CX), while ours replaces the first (XC). However, the number of replacements is the same in both cases.

There are several promising directions for future work that aim to further enhance the performance of `RLZ-RePair`. One is the choice of reference, which plays an important role in the performance of our method. In our experiments, we randomly selected the sequences that made up the reference; however, some sequences in the reference likely contributed little to improving the RLZ parse. Choosing the reference in a more systematic way like described in [3] may lead to significantly better performance. Another promising approach we recently implemented involves making non-explicit phrases that span only a small interval of the reference explicit from the start. Preliminary experiments show that this reduces the number of phrases, but increases memory usage, as more characters are made explicit that might otherwise have been kept compressed. With further refinement, this approach could help keep resource usage low, even with a suboptimal reference.

## A RLZ-RePair: Step-by-Step Example

Suppose we want to RLZ-RePair the following text (*T*)

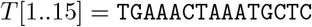

and we choose the following reference (*R*)

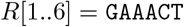

then the Relative Lempel-Ziv (RLZ) parse of *T* with respect to *R* is

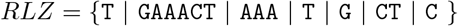

which can be represented as (*p*_*i*_, *ℓ*_*i*_) pairs

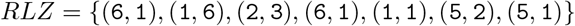

where *p*_*i*_ is the starting position of the match in *R*, and *ℓ*_*i*_ is the match length. For convenience, we reinterpret each (*p*_*i*_, *ℓ*_*i*_) pair to mark the start (*s*_*i*_) and end (*e*_*i*_) of its corresponding interval in *R*, as follows

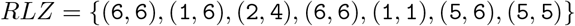

where *s*_*i*_ = *p*_*i*_ and *e*_*i*_ = *p*_*i*_ + *ℓ*_*i*_ − 1. These RLZ phrases are referred to as **non-explicit** phrases. Figure A1 provides a visual summary of the key information derived from the RLZ parse prior to the start of RePair compression.

**Figure A1.**
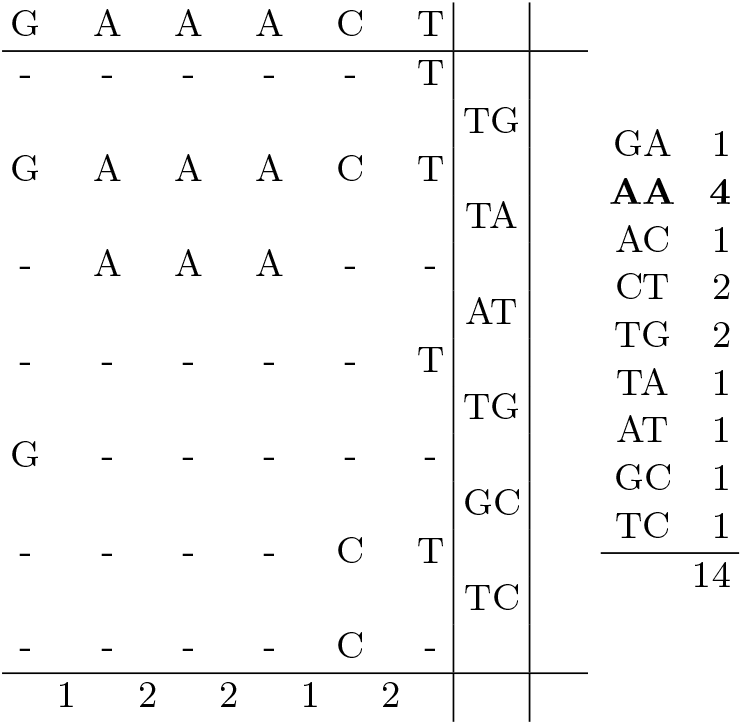
Visual representation of the RLZ parse and the derived bigram frequencies prior to RePair compression. The reference text **(left, top row)** and non-explicit phrases **(left, middle rows)** are shown, with non-explicit phrases positioned to illustrate their overlap with the reference. Bigrams crossing phrase boundaries are shown **(left, right column)**. Total bigram frequencies in *T* are shown **(right)**, with the most frequent bigram bolded. Total bigram frequencies are computed by summing those within non-explicit phrases **(left, bottom row)** and those crossing phrase boundaries. Notice the total frequency is 1 less than the number of characters in *T* .

The RePair algorithm iteratively replaces the most frequent bigram with a new non-terminal symbol until no bigram occurs more than once. In Figure A1, the frequency table shows that the most frequent bigram is AA. Before performing a replacement, RLZ-RePair must ensure that no non-explicit phrase is invalidated by checking phrase and source boundary conditions. As shown in Figure A1, `AA` neither crosses phrase boundaries nor is partially overlapped by any phrase in the reference, making it safe to replace. Since all occurrences of `AA` in the text are fully contained within non-explicit phrases, replacements can be carried out in the text by proxy through substitutions in the reference. Although `AA` appears twice in the reference, its consecutive occurrences means only one replacement is necessary. We replace all occurrences of `AA` in the reference with a new symbol `4`. Figure A2 shows the resulting information after the replacement. If the non-explicit phrases are long, then it expected that most replacements occurring in the text can be resolved by simply doing the replacements in the reference.

**Figure A2.**
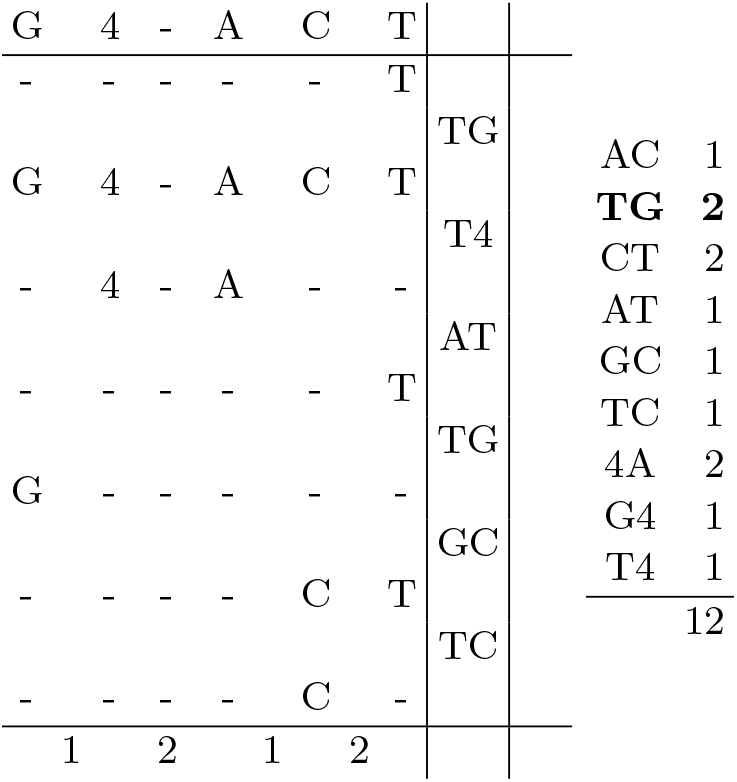
Visual representation of the RLZ parse and the derived bigram frequencies after replacing AA with 4. The reference text **(left, top row)** and non-explicit phrases **(left, middle rows)** are shown, with non-explicit phrases positioned to illustrate their overlap with the reference. Bigrams crossing phrase boundaries are shown **(left, right column)**. Total bigram frequencies in *T* are shown **(right)**, with the most frequent bigram bolded. Total bigram frequencies are computed by summing those within non-explicit phrases **(left, bottom row)** and those crossing phrase boundaries. Notice the total frequency is 1 less than the number of characters in *T* .

Figure A2 shows a tie among the most frequent bigrams remaining in the text: `TG, CT`, and `4A`, each with a frequency of 2. We choose to replace `TG`. Before the replacement, we must again check for phrase or source boundary conditions that could invalidate the non-explicit phrases. We find that a phrase boundary violation exists. Since both occurrences of `TG` span consecutive phrases and are not fully contained within any single phrase, the replacement cannot be applied directly to the reference like we did previously. To resolve this issue, we make the characters **explicit**, i.e. literals, by partially decompressing the ends of the affected phrases and storing them in plain text, as shown in Figure A3. Once the necessary characters are made explicit, the replacement can safely be applied to the reference and the explicit phrases. In this case, the bigram `TG` does not occur in the reference, so all the replacements occur in the explicit phrases. We replace all occurrences of `TG` in the explicit phrases with a new symbol 5. Figure A4 shows the resulting information after the replacement.

Figure A4 shows another tie among the most frequent bigrams remaining in the text: `CT` and `4A`, each with a frequency of 2. We choose to replace `CT`. Again, we check to see if there are any phrase or source boundary conditions prior to the replacement. We observe no occurrences of `CT` crossing phrase boundaries, so no phrase boundary violations need to be addressed. However, we do observe a non-explicit phrase partially overlapping an occurrence of `CT` in the reference. Therefore, a source boundary violation exists. To resolve this, we shorten the non-explicit phrase by removing its end character and making it explicit, like we did previously to resolve the phrase boundary condition, as shown in Figure A5. After making the necessary characters explicit, we can safely do the replacement in the reference and explicit phrases. Since both occurrences of the bigram `CT` in the text are fully covered by non-explicit phrases, the replacement only has to be done in the reference. We replace all occurrences of `CT` in the reference with a new symbol 6. Figure A6 shows the resulting information after the replacement.

**Figure A3.**
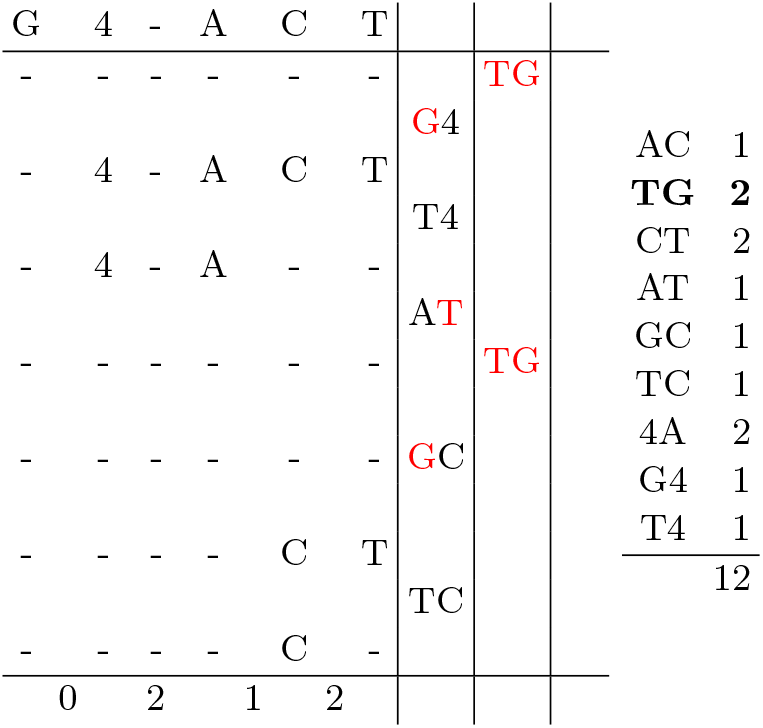
Visual representation of the RLZ parse and the derived bigram frequencies after resolving phrase and source boundary conditions for the bigram `TG` prior to its replacement. The reference text **(left, top row)** and non-explicit phrases **(left, middle rows)** are shown, with non-explicit phrases positioned to illustrate their overlap with the reference. Bigrams crossing phrase boundaries are shown **(left, middle column)**. Explicit phrases are shown **(left, right column)**. Explicit characters are colored red. Total bigram frequencies in *T* are shown **(right)**, with the most frequent bigram bolded. Total bigram frequencies are computed by summing those within non-explicit phrases **(left, bottom row)**, those crossing phrase boundaries, and those within explicit phrases. Notice the total frequency is 1 less than the number of characters in *T* .

**Figure A4.**
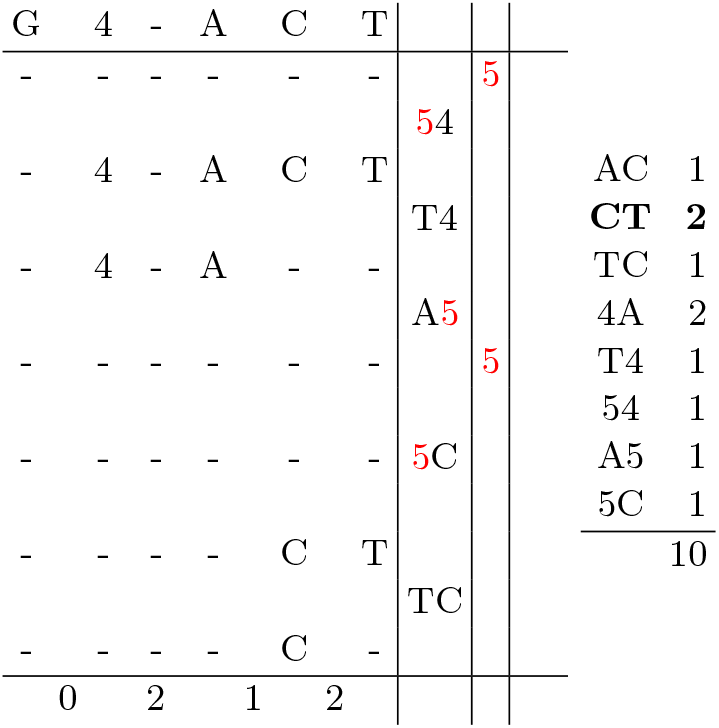
Visual representation of the RLZ parse and the derived bigram frequencies after replacing `TG` with `5`. The reference text **(left, top row)** and non-explicit phrases **(left, middle rows)** are shown, with non-explicit phrases positioned to illustrate their overlap with the reference. Bigrams crossing phrase boundaries are shown **(left, middle column)**. Explicit phrases are shown **(left, right column)**. Explicit characters are colored red. Total bigram frequencies in *T* are shown **(right)**, with the most frequent bigram bolded. Total bigram frequencies are computed by summing those within non-explicit phrases **(left, bottom row)**, those crossing phrase boundaries, and those within explicit phrases. Notice the total frequency is 1 less than the number of characters in *T* .

**Figure A5.**
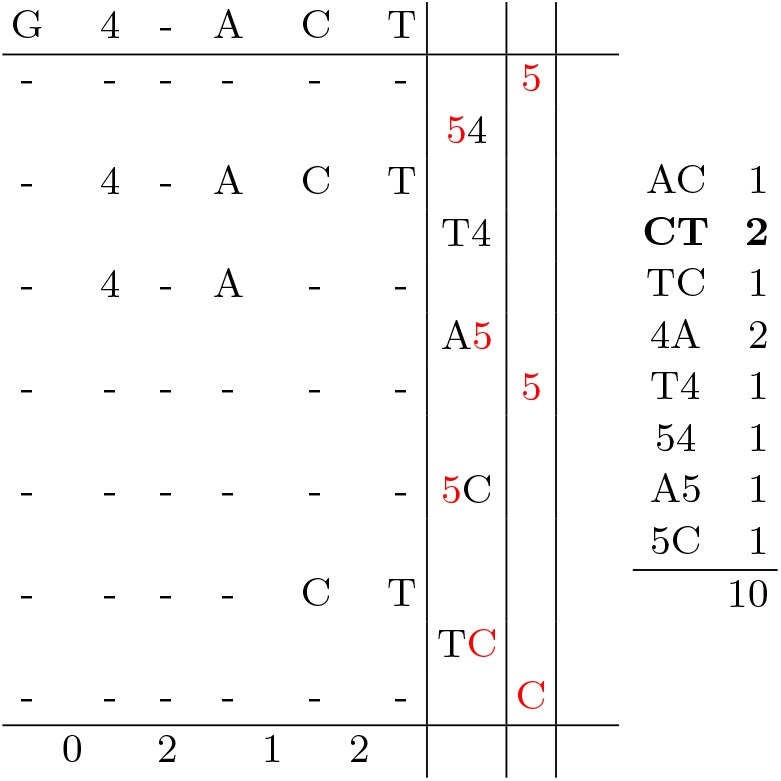
Visual representation of the RLZ parse and the derived bigram frequencies after resolving phrase and source boundary conditions for the bigram `CT` prior to its replacement. The reference text **(left, top row)** and non-explicit phrases **(left, middle rows)** are shown, with non-explicit phrases positioned to illustrate their overlap with the reference. Bigrams crossing phrase boundaries are shown **(left, middle column)**. Explicit phrases are shown **(left, right column)**. Explicit characters are colored red. Total bigram frequencies in *T* are shown **(right)**, with the most frequent bigram bolded. Total bigram frequencies are computed by summing those within non-explicit phrases **(left, bottom row)**, those crossing phrase boundaries, and those within explicit phrases. Notice the total frequency is 1 less than the number of characters in *T* .

Figure A6 shows that the most frequent bigram is `4A`. Luckily, there are no phrase or source boundary violations with this bigram. Since it does not occur in any explicit phrase, we can directly do the replacement in the reference to cover all the replacements in the text. We replace all occurrences of `4A` in the reference with a new symbol 7. Figure A7 shows the resulting information after the replacement.

We see from Figure A7 that the frequency of all bigrams in the text is 1, which indicates that the algorithm terminates. We can then write the compressed (C) sequence and rule (R) files to disk. The remaining non-explicit phrases at this point can be decompressed by using the reference. If the text is repetitive and the reference captures its repetitiveness, then a large percentage of the remaining characters should still be a part of the non-explicit phrases. The resulting C and R files for the example is as follows:

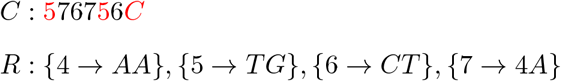

**Figure A6.**
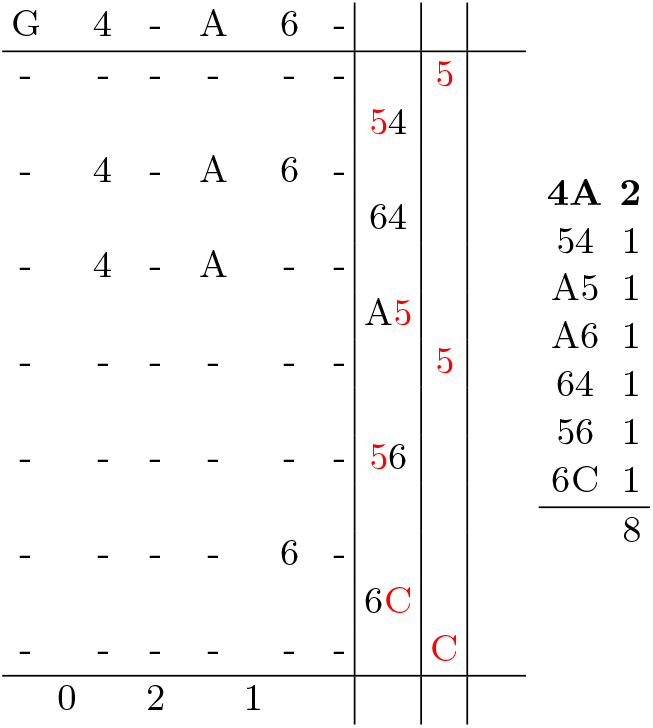
Visual representation of the RLZ parse and the derived bigram frequencies after replacing `CT` with `6`. The reference text **(left, top row)** and non-explicit phrases **(left, middle rows)** are shown, with non-explicit phrases positioned to illustrate their overlap with the reference. Bigrams crossing phrase boundaries are shown **(left, middle column)**. Explicit phrases are shown **(left, right column)**. Explicit characters are colored red. Total bigram frequencies in *T* are shown **(right)**, with the most frequent bigram bolded. Total bigram frequencies are computed by summing those within non-explicit phrases **(left, bottom row)**, those crossing phrase boundaries, and those within explicit phrases. Notice the total frequency is 1 less than the number of characters in *T* .

**Figure A7.**
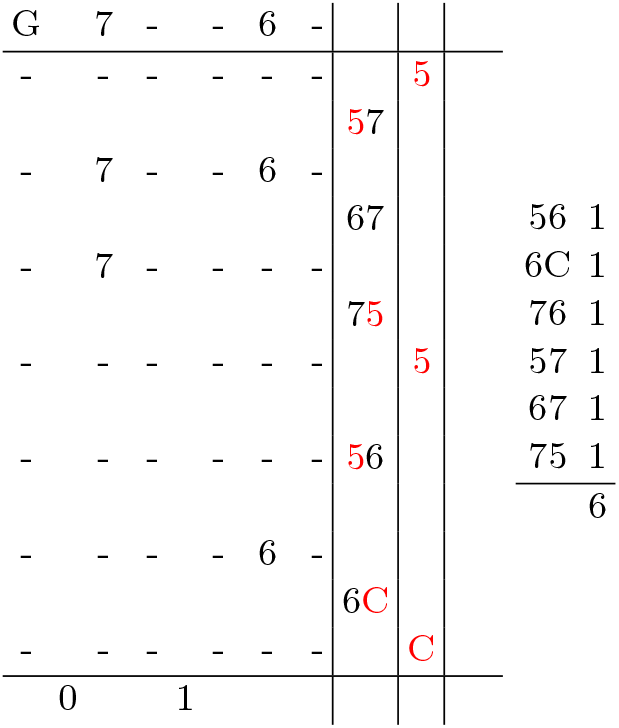
Visual representation of the RLZ parse and the derived bigram frequencies after replacing `4A` with `7`. The reference text **(left, top row)** and non-explicit phrases **(left, middle rows)** are shown, with non-explicit phrases positioned to illustrate their overlap with the reference. Bigrams crossing phrase boundaries are shown **(left, middle column)**. Explicit phrases are shown **(left, right column)**. Explicit characters are colored red. Total bigram frequencies in *T* are shown **(right)**, with the most frequent bigram bolded. Total bigram frequencies are computed by summing those within non-explicit phrases **(left, bottom row)**, those crossing phrase boundaries, and those within explicit phrases. Notice the total frequency is 1 less than the number of characters in *T* .

## References

1 Djamal Belazzougui, Patrick Hagge Cording, Simon J Puglisi, and Yasuo Tabei. Access, rank, and select in grammar-compressed strings. In the Proceedings of the 23rd Annual European Symposium (ESA), pages 142–154. Springer, 2015.

2 Philip Bille, Inge Li Gørtz, and Nicola Prezza. Practical and effective Re-Pair compression. arXiv preprint 1704.08558, 2017.

3 Philip Bille, Inge Li Gørtz, Simon J. Puglisi, and Simon R. Tarnow. Hierarchical Relative Lempel-Ziv Compression. In the Proceedings of the 21st International Symposium on Experimental Algorithms (SEA), volume 265, pages 18:1–18:16, 2023.

4 Philip Bille, Gad M Landau, Rajeev Raman, Kunihiko Sadakane, Srinivasa Rao Satti, and Oren Weimann. Random access to grammar-compressed strings and trees. SIAM Journal on Computing, 44(3):513–539, 2015.

5 Francisco Claude, Antonio Farina, Miguel A Martínez-Prieto, and Gonzalo Navarro. Compressed q-gram indexing for highly repetitive biological sequences. In the Proceedings of the 10th IEEE International Conference on BioInformatics and BioEngineering (BIBE), pages 86–91. IEEE, 2010.

6 Francisco Claude and Gonzalo Navarro. Fast and compact web graph representations. ACM Transactions on the Web, 4(4):1–31, 2010.

7 Thomas H. Cormen, Charles E. Leiserson, Ronald L. Rivest, and Clifford Stein. Introduction to Algorithms, 3rd Edition. MIT Press, 2009.

8 Diego Alejandro Díaz Domínguez. Data structures and algorithms for analyzing DNA sequences in compressed space, 2021.

9 Travis Gagie, Tomohiro I, Giovanni Manzini, Gonzalo Navarro, Hiroshi Sakamoto, and Yoshimasa Takabatake. Rpair: Rescaling RePair with Rsync. In the Proceedings of the 26th International International Symposium on String Processing and Information Retrieval (SPIRE), pages 35–44, 2019.

10 Matthias Gallé. Investigating the effectiveness of BPE: The power of shorter sequences. In Proceedings of the Conference on Empirical Methods in Natural Language Processing and the 9th International Joint Conference on Natural Language Processing (EMNLP-IJCNLP), pages 1375–1381, November 2019.

11 Rodrigo González and Gonzalo Navarro. Compressed text indexes with fast locate. In the Proceedings of the 33rd Annual Symposium on Combinatorial Pattern Matching (CPM), pages 216–227. Springer, 2007.

12 Ben Hu, Hua Guo, Peng Zhou, and Zheng-Li Shi. Characteristics of SARS-CoV-2 and COVID-19. Nature Reviews Microbiology, 19(3):141–154, 2021.

13 Dominik Kempa and Tomasz Kociumaka. Lempel-Ziv (LZ77) factorization in sublinear time. In the Proceedings of the IEEE 65th Annual Symposium on Foundations of Computer Science (FOCS), pages 2045–2055. IEEE, 2024.

14 Justin Kim, Rahul Varki, Marco Oliva, and Christina Boucher. Re2Pair: Increasing the Scalability of RePair by Decreasing Memory Usage. In the Proceedings of the 32nd Annual European Symposium on Algorithms (ESA), volume 308, pages 78:1–78:15, 2024.

15 László Kozma and Johannes Voderholzer. Theoretical analysis of byte-pair encoding. arXiv preprint 2411.08671, 2024.

16 Shanika Kuruppu, Simon J Puglisi, and Justin Zobel. Relative Lempel-Ziv compression of genomes for large-scale storage and retrieval. In the Proceedings of the 17th International Symposium on String Processing and Information Retrieval (SPIRE), pages 201–206, 2010.

17 N Jesper Larsson and Alistair Moffat. Off-line dictionary-based compression. Proceedings of the IEEE, 88(11):1722–1732, 2000.

18 Heng Li. seqtk: A fast and lightweight tool for processing fasta/fastq sequences, version 1.3-r106. https://github.com/lh3/seqtk, 2013. Version 1.3-r106, Accessed: 2025-04-22.

19 Heng Li, Xiaoping Hong, Liping Ding, Shuhui Meng, Rui Liao, Zhenyou Jiang, and Dongzhou Liu. Sequence similarity of SARS-CoV-2 and humans: Implications for SARS-CoV-2 detection. Frontiers in Genetics, 13:946359, 2022.

20 Heng Li and Jiazhen Rong. Bedtk: finding interval overlap with implicit interval tree. Bioinformatics, 37(9):1315–1316, 2021.

21 Markus Lohrey, Sebastian Maneth, and Roy Mennicke. XML tree structure compression using RePair. Information Systems, 38(8):1150–1167, 2013.

22 Takuya Mieno, Shunsuke Inenaga, and Takashi Horiyama. RePair Grammars Are the Smallest Grammars for Fibonacci Words. In the Proceedings of the 33rd Annual Symposium on Combinatorial Pattern Matching (CPM), pages 26:1–26:17, 2022.

23 Felix Mölder, Kim Philipp Jablonski, Brice Letcher, Michael B Hall, Christopher H Tomkins-Tinch, Vanessa Sochat, Jan Forster, Soohyun Lee, Sven O Twardziok, Alexander Kanitz, Andreas Wilm, Manuel Holtgrewe, Sven Rahmann, Sven Nahnsen, and Johannes Köster. Sustainable data analysis with Snakemake. F1000Research, 10, 2021.

24 Gonzalo Navarro, Víctor Sepúlveda, Mauricio Marín, and Senén González. Compressed filesystem for managing large genome collections. Bioinformatics, 35(20):4120–4128, 2019.

25 Craig G Nevill-Manning and Ian H Witten. Identifying hierarchical structure in sequences: A linear-time algorithm. Journal of Artificial Intelligence Research, 7:67–82, 1997.

26 Ivan Provilkov, Dmitrii Emelianenko, and Elena Voita. BPE-dropout: Simple and effective subword regularization. In the Proceedings of the 58th Annual Meeting of the Association for Computational Linguistics (ACL), pages 1882–1892, 2020.

27 The 1000 Genomes Project Consortium. A global reference for human genetic variation. Nature, 526:68–74, 2015.

